# Analyses of single extracellular vesicles from non-small lung cancer cells to reveal effects by Epidermal growth factor inhibitor treatments

**DOI:** 10.1101/2022.10.14.512252

**Authors:** Fredrik Stridfeldt, Sara Cavallaro, Petra Hååg, Rolf Lewensohn, Jan Linnros, Kristina Viktorsson, Apurba Dev

## Abstract

Precision cancer medicine have changed the treatment landscape of non-small cell lung cancer (NSCLC) as illustrated by tyrosine kinase inhibitors (TKIs) towards mutated Epidermal growth factor receptor (EGFR). Yet, responses to such TKIs e.g., erlotinib and osimertinib among patients are heterogenous and there is a need for non-invasive blood-based analytics to follow treatment response and reveal resistance to improve patient’s treatment outcome. Recently, extracellular vesicles (EVs) have been identified as an important source of tumor biomarkers promising to revolutionize liquid biopsy-based diagnosis of cancer. However, high heterogeneity has been a major bottleneck. The pathological signature is often hidden in the differential expression of membrane proteins in a subset of EVs which are difficult to identify with bulk techniques. Using a fluorescence-based approach, we for the first time demonstrate that the single-EV technique can be used to monitor the treatment response of targeted cancer therapies such as TKIs towards EGFR. To test the hypothesis, we analyzed the membrane proteins of native EVs extracted from EGFR-mutant NSCLC cell line, both prior and post treatment with EGFR-TKIs erlotinib or osimertinib. The selected cell line being refractory to erlotinib and responsive to osimertinib makes it a suitable model system. The expression level of five surface proteins; two common tetraspanins (CD9, CD81) and three markers of specific interest in lung cancer (EGFR, PD-L1, HER2) were studied. The data suggest that in contrast to erlotinib, the osimertinib treatment increases the population of PD-L1, EGFR and HER2 positive EVs while the expression level per EV decreases for all the three markers. The PD-L1 and HER2 expressing EV population seems to increase by several fold because of osimertinib treatment. The observations agree with the previous reports performed on cellular level indicating the biomarker potential of EVs for liquid-biopsy based monitoring of targeted cancer treatments.

**Highlights:**

- Membrane protein analyses of single EVs may reveal distinct differences when lung cancer cells are refractory vs responsive under different EGFR-TKI treatments.
- Comparison of 1^st^ generation erlotinib and 3^rd^ generation osimertinib shows clear signature on the expression of PD-L1, EGFR, HER2 on single EVs
- Colocalization showed a change in common marker combinations before after treatment.
- PD-L1 expression per vesicle decreases while the number of PD-L1 positive EVs increases as a result of osimertinib treatment, indicating that such signature may not be detectable under bulk analysis

## Introduction

Extracellular vesicles (EVs) are nanoscale lipid-enclosed vesicles released by all cells. For a long time, the scientific community overlooked the importance of EVs since they were only assumed to participate in the control of cellular homeostasis, i.e., to function as “waste carriers” to get rid of unneeded materials [1]. However, after discovering EVs ability to transfer active molecules such as RNA and proteins, EVs have gained strong interest as a potential key player in different fields such as physiology, pathology including cancer, neurodegenerative disorders, and drug delivery strategies, to mention a few [2], [3], [4]. EVs are broadly classified based on their size and route of biogenesis, such as exosomes which originate from the endolysosomal pathway; microvesicles which bud from the plasma membrane and apoptotic bodies which are released from a dying cell [5]. Due to the difficulties in their selective isolation, EVs are often classified as small EVs (sEVs; 100-200 nm) and medium/large EVs (m/lEVs; >200 nm) [6]. However, it is increasingly evident that there exist subclasses of EVs in each group making them phenotypically and functionally more heterogeneous [6]. Identifying subpopulations of EVs is currently one of the major efforts in the field [7].

The treatment regimen for non-small cell lung cancer (NSCLC) patients with advanced or metastatic disease have changed during the last decade where molecular profiling have identified oncogenic mutations e.g. *epidermal growth factor receptor (EGFR), Kirsten rat sarcoma virus (KRAS, G12C* mutations) and *v-raf murine sarcoma viral oncogene homolog B1 (BRAF)* or gene fusions/rearrangements, that have allowed precision cancer medicine (PCM) treatments using receptor tyrosine kinase (RTK) inhibitors (TKIs) [8] [9] [10]. Some NSCLC patients have also benefited from re-activation of the immune system towards the tumor using so-called immune checkpoint inhibitors (ICI). About 12-30% of all NSCLC is due to mutated *EGFR*, which are treated with EGFR-TKIs. These include 1^st^ generation gefitinib and erlotinib but lately also with 2^nd^ generations TKIs i.e., afatinib, dacomitinib and importantly also by the 3^rd^ generation osimertinib. EGFR-TKIs may for some patients allow a long progression-free survival, with an osimertinib treatment showing the longest progression-free survival [9]. However, all patients relapse calling for ways to early monitoring of treatment response. It is well established that compensatory genomic alterations in EGFR, e.g., mutations in kinase pocket is a *bona fide* resistance mechanism for EGFR-TKIs while the genomic changes associated to 1^st^ and 3^rd^ generation EGFR-TKIs differ in this respect [11]. Moreover, for a subset of the EGFR-TKI resistant cases, the resistance is attributed to alteration in so-called by-pass signaling drivers which in context of 1^st^ generation includes among others HER2, MET, IGF-1R, AXL and EPHA2 [11]. Thus, treatment monitoring analytics for EGFR-mutant NSCLC patients such as those based on EVs from plasma or serum samples during treatment should also include ways to follow these signaling cascades alongside cell free DNA in plasma.

Profiling of EVs for early NSLCC diagnostics using bulk EVs samples for protein or RNA analyses have deciphered certain markers [12] [13] [14] [15]. Multiple studies in different tumor types have also demonstrated that EGFR signaling may influence the EVs released in terms of tetraspanin CD9, CD63, CD81 expression as well as other proteins [16] [17]. Moreover, it has been demonstrated that EVs contain various immune signaling components including PD-L1 which could contribute to an immune suppressive phenotype seen among NSCLC cases and which influence immune checkpoint inhibitor treatments [18] [19]. Yet, most of these EV studies have focused on bulk analyses of EVs, thereby missing the hidden signatures in the heterogenous EV population as well as the aspect of tumor heterogeneity which can have major impact on the diagnostic accuracy and scope.

Using a fluorescence-based single-EV approach in this study, we demonstrate that the membrane proteins of EVs reveals interesting differences which can potentially be used for early detection of treatment response of targeted cancer therapies using liquid biopsy. For this purpose, we analyzed five membrane proteins (CD9, CD81, EGFR, HER2 and PD-L1) on individual EVs extracted from an EGFR-mutant NSCLC cell line (H1975), both prior and post treatment with EGFR-TKIs erlotinib or osimertinib. The selection of the treatment, i.e. 1^st^ generation erlotinib and 3^rd^ generation osimertinib, is motivated to highlight the differences when the cell is refractory vs responsive to the applied drugs. To further increase our understanding, we compared the results with the non-EGFR-TKI medication cisplatin. The results indicate in contrast to erlotinib and cisplatin, osimertinib treatment increases the HER2, EGFR and PD-L1 positive EV population. However, the average expression of the same three proteins on individual EVs were found to drop as result of the osimertinib treatment. Interesting differences were also observed in the distribution of expression levels on individual EVs positive for either of HER2, EGFR and PD-L1, indicating possibilities to use such signatures to follow treatment response of targeted cancer therapy in liquid biopsy-based analysis.

## Materials and methods

### Reagents

High purity deionized water (DIW) with a resistivity of 18 MΩ cm was used throughout all the experiments. Casein (C5890) in powder, Phosphate-buffered saline (PBS, P4417) in tablets and Poly-L-Lysine (PLL, P2636) were purchased from Sigma-Aldrich (Burlington, MA, United States). Anti-CD9 antibody conjugated with VioBlue (#130-118-809) was purchased from Miltenyi Biotec (Bergisch Gladbach, NRW, Germany), while anti-EGFR R-PE (#1P-680-T100), anti-Human CD274/PD-L1 APC (#1A-177-T100), anti-Human CD340 / HER2 APC (#1A-832-T100) were purchased from Exbio (Vestec, CBR, Czech Republic). Anti-CD81-APC (A87789) was purchased from Beckman Coulter (Brea, CA, United States). All the antibodies used were monoclonal.

### Fluorescent (FL) measurement set up

The fluorescent (FL) measurements of the vesicles were performed with an inverted microscope from Zeiss (Axio Observer 7; Oberkochen, BW, Germany). The microscope was equipped with a Colibri 5 containing four light-emitting diode (LED) modules for exciting the samples. The three modules used in this paper were centered around 385 nm, 555 nm, and 630 nm, respectively. The 385 nm channel was used for exciting CD9-VioBlue, the 555 nm channel for EGFR-rPE, and 630 nm channel for PD-L1/HER2/CD81-APC. Each LED module held a suitable excitation filter. All measurements were performed with a 100× oil immersion objective lens.

The images (captured area of each image: 133.12 μm × 133.12 μm) were taken using a Hamamatsu CCD Camera (Orca Flash 4) with the ZEN 3.0 program on a computer connected to the microscope. The images were captured in the three excitation channels for 2 seconds each. All LEDs were used at maximum capacity except for the one centered around 385 nm, which was limited to 60% to prevent increased background noise and quick photobleaching of fluorophore signal. The particles were identified and quantified by the ZEN 3.0 program. Both spatial and intensity thresholds were used in this process allowing outliers in the samples and protein profiles to be revealed. These thresholds were based on control images and tailored to exclude background noise as well as agglomerates of fluorophore molecules and dust. For comparative reasons, similar thresholds were used for all experiments. Background correction was carried out based on the rolling ball algorithm with a 20-pixel radius. After analysis each spot within a defined range of area and intensity was circled. The intensity of a vesicle was then calculated as an average of the pixel intensities making up each spot.

### Cell treatment and isolation of extracellular vesicles

In this work EVs was obtained from the NSCLC H1975 cell culture media. The H1975 cell line (ATCC^®^ CRL-5908^TM^, LGC Standards, Teddington Middlesex, United Kingdom) is a commonly used *in vitro* model for epidermal growth factor receptor (EGFR) -driven NSCLC. The H1975 cells are known to be refractory to 1^st^ generation EGFR-TKI, e.g., erlotinib as they alongside the EGFR sensitizing mutation, exon 21, L858R, also have T790M in exon 20, a gatekeeper mutation that prevent efficacy of 1st generation EGFR-TKI [20]. Importantly, H1975 cells are sensitive to 3^rd^ generation osimertinib which also was applied in this study [21]. For collecting EVs from cell culture media, H1975 cells (3 ×10^6^ 175 cm^2^ flask) were treated with either erlotinib (1 μM) or osimertinib (0.1 μM) for about 48h with untreated cells used in parallel. In one set of experiments, H1975 cells were also treated with cisplatin (12.5 or 25 μM (Cisplatin Ebewe, Apoteket Sverige AB, Stockholm, Sweden)) for 48h. All cells were seeded in RPMI media (supplemented with 10% fetal bovine serum (FBS) and 2 mM L-glutamine (Gibco, Life Technologies, Stockholm, Sweden) and were for 48h prior experiments cultured in RPMI-media with exosome-depleted FBS (10%, Gibco™ A2720801, Fisher Scientific, Gothenburg, Sweden). The EVs were isolated from about 75 ml cell culture media/sample which was collected and cleared from cell debris by centrifugation first for 5 minutes at 200 RCF and thereafter for 20 minutes at 720 RCF for 20 minutes. The EVs were thereafter captured from the supernatant by size exclusion chromatography (SEC) on the qEVoriginal Columns (Izon Science, Oxford, United Kingdom) giving EVs with size from 30-300 nm. The detailed description of EV our isolation procedure is reported in Stiller *et al* [22].

The particle size and particle/ml in the samples were analyzed using the ZetaView (Particle Metrix GmbH, Ammersee, Germany). The samples were diluted in PBS to give appropriate particles/frame. For all samples the following settings were used: cell temperature: 25°C; camera sensitivity for all samples: 80; shutter: 100; number of cycles: 3.

Capturing and protein profiling platform for single extracellular vesicles

A detailed description of the fluorescence based single EV platform and optimization of the method is explained in our previous report [23]. A schematic of the experimental procedure is presented in Figure 1. After harvesting, EVs were first characterized with standard EV characterization methods which includes NTA and western blot. For fluorescence imaging, EVs were unbiasedly captured on a gridded coverslip (Ibidi 10817, Gräfelfing, Munich, Germany). The glass coverslip was cleaned with a 5:1:1 solution of DIW, H2O2, and NH4OH (88°, 10 minutes) and activated with Poly-L-Lysine (PLL, 0.01% w/v in DIW for 30 minutes). A four-well insert (Ibidi 80469, Gräfelfing, Munich, Germany) was attached to the slip enabling up to three simultaneous antibody-probing experiments and one control environment without any vesicles but with antibodies. The slip was bathed in DIW for 1 hour prior attaching the insert to ease the binding. The EV samples were diluted to a final concentration of 2×10^8^ particles/ml using 1×PBS before they were adhered to the coverslip. The vesicles were incubated on the substrates for one hour at room temperature during which time the vesicle adhered electrostatically to the PLL. To ensure likewise number of particles incubated across each experiment, the EV sample incubation was performed with the same total number of particles adjusted according to NTA data. To reduce non-specific binding between the antibodies and the surface, the substrate was incubated overnight with casein as blocking agent. To further avoid nonspecific binding, the fluorophore-conjugated antibodies were to interact with the proteins in a casein environment. The antibodies were diluted to a concentration of 4 nM before incubation. The concentration was chosen to be higher than the concentration of the vesicles (with a >1:1 200 EV:antibody ratio) to avoid the number of antibodies being a limiting factor in the particle identification.

**Figure 1.**
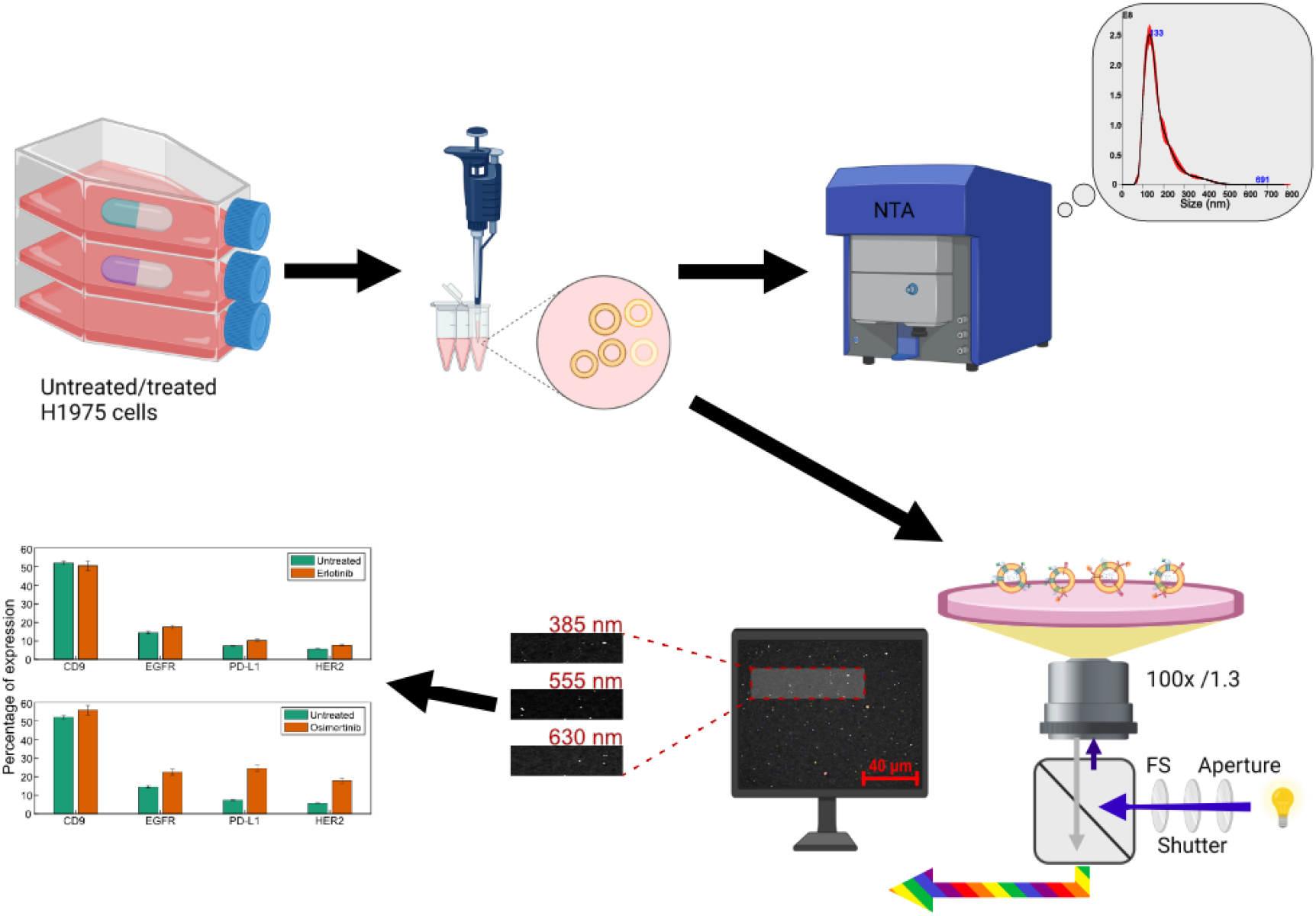
Schematic representation of the experimental procedure. EVs are collected from the NSCLC H1975 cell culture media. One sample was collected without any treatment, and four other samples where the cells were either treated with erlotinib (1 μM), osimertinib (0.1 μM), or cisplatin (12.5 μM and 25 μM) for 48 h before harvest. The size and concentration of the samples were profiled using a nanoparticle tracking analysis (NTA). Along with the profiling, a coverslip coated with Poly-L-Lysine (PLL) to bind EVs and casein to reduce unspecific binding of when probing with antibodies were used. The glass slide was probed with EVs and subsequently incubated with monoclonal antibody-fluorophore conjugates targeting a specific surface protein. In an inverted microscope, the sample was exposed to LED lights of specific wavelengths where excited fuorophore signal indicates binding of the antibody to a protein in individual EVs. The results were captured with a CCD camera for analysis. Figure partly created using BioRender.com.

## Results

### Characterization of H1975 cells and EVs

The cell viability of the H1975 cell line after exposure to the different treatments is presented in Figure 2A. Cells were counted with trypan blue and viable cells (not blue) were analyzed. Representative images of the cell morphology of the samples are presented in Figure S1. The results agree with our earlier reports that osimertinib at this concentration causes a 50% reduction in cell viability while with erlotinib only minor effect on cell viability is seen [24]. In accordance with the Minimum Information for Studies of Extracellular Vesicles 2018 (MISEV2018) guidelines [6], the presence of proteins in the vesicles was confirmed by western blot analysis of EGFR, CD73, TSG101, and CD9, with GADPH being a negative marker. PARP was included to show indication of apoptosis in the cell samples.

**Figure 2.**
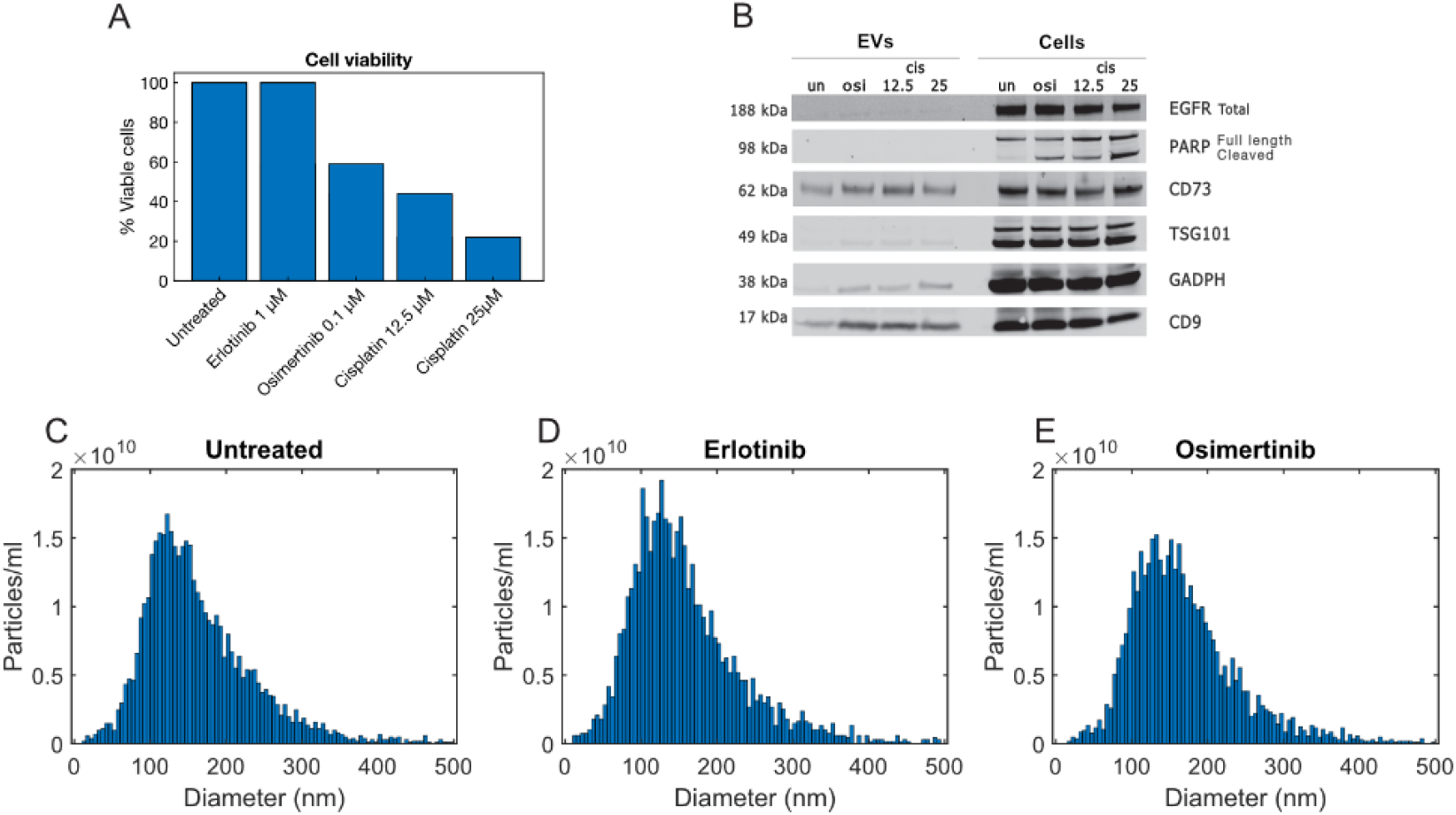
Characterization of cells and EVs. Cell viability for untreated, erlotinib (1 μM) osimertinib (0.1 μM) and cisplatin (12.5 μM and 25 μM) cells. Viability of erlotinib from an earlier investigation, see [24]. Cells were treated 48 h before counting. (A). Expression of EGFR, PARP, CD73, TSG101, GADPH, and CD9 in the different H1975 cell and EV samples as measured by western blotting (B). Size and concentration analysis using ZetaView for Untreated (C), erlotinib treated (D), and osimertinib treated (E) samples used in the treatment study.

The resulting western blots are shown in Figure 2B. Earlier western blot profiling of EVs from these treatments have confirmed expression of CD9 while little or no contamination with cellular proteins [24] [22]. Nanoparticle tracking analyses of the untreated, osimertinib, and erlotinib samples have revealed that isolated EVs were of EVs size and the median sizes of about 145 nm, see Figure 2C-E.

We next went on and studied the expression of CD9 and CD81 as well as some targets of relevance to EGFR-TKI response and resistance in *EGFR*-mutant NSCLC i.e., EGFR and HER2 on individual EVs. Further, we included the immune checkpoint protein PD-L1 which earlier have been reported in EGFR-mutant NSCLC cells [25]. Before imaging with EVs, the background signal from the substrate was thoroughly investigated to ensure high signal to noise ratio and low non-specific signals. Figure S2 shows the intensity counts for each channel measured on a coverslip with identical chemical functionalization but without any EVs adhered to it. No difference could be observed on the overall background intensities between the surfaces without (Figure S2A) and with antibodies (Figure S2B). However, depending on the antibodies few discrete spots could be observed per imaging area as shown in Figure S3. The number was deemed low enough to not have a severe impact on the results.

Earlier report have shown procedures to study many more surface proteins on the same sample using longer steps of photobleaching [26]. However, due to the limiting surface area of EVs, we anticipate that steric hindrance may start to dominate impeding the accuracy of the measurement. Therefore, we measured up to a maximum of three markers at a time and repeated the experiments twice with different combination of antibodies. CD9 and EGFR were investigated in all the experiments the third target was alternated between PD-L1, HER2, and CD81. Having CD9 and EGFR as a common marker, allowed us to compare the results among different experiments. For each sample and experiment, 15 microscope images were considered for further studies. Figure 3A shows the number of EVs positive for the two tetraspanins CD9 and/or CD81 in the untreated and the EGFR-TKI treated samples. Even though the samples were diluted to a shared concentration of 2**×**10^8^ particles/ml, according to NTA data, a big difference in the number of tetraspanins positive EVs was identified in the fluorescent measurement. The number of EVs positive for each tetraspanin in the erlotinib sample was reduced by 80% ± 5% of the untreated number. The osimertinib sample shows a similar trend, with drops of approximately 80% from the untreated to the treated samples. The drop may indicate to a potential decrease in the overall EV count not detected by NTA. As a result, for further analysis, we decided to normalize the data with respect to CD9 positive EVs for each sample. This way, we could compare the samples even if the EV counts differed. The percentage of EVs normalized to the number of CD9-positive EVs in each of the tested samples is presented as bar plot in Figure 3B. Normalization to CD81 instead of CD9 (Figure S4) shows similar behavior and justifies this standardization. Both CD9 and CD81 are common markers for EVs and were deemed suitable as the roll of normalization protein [27].

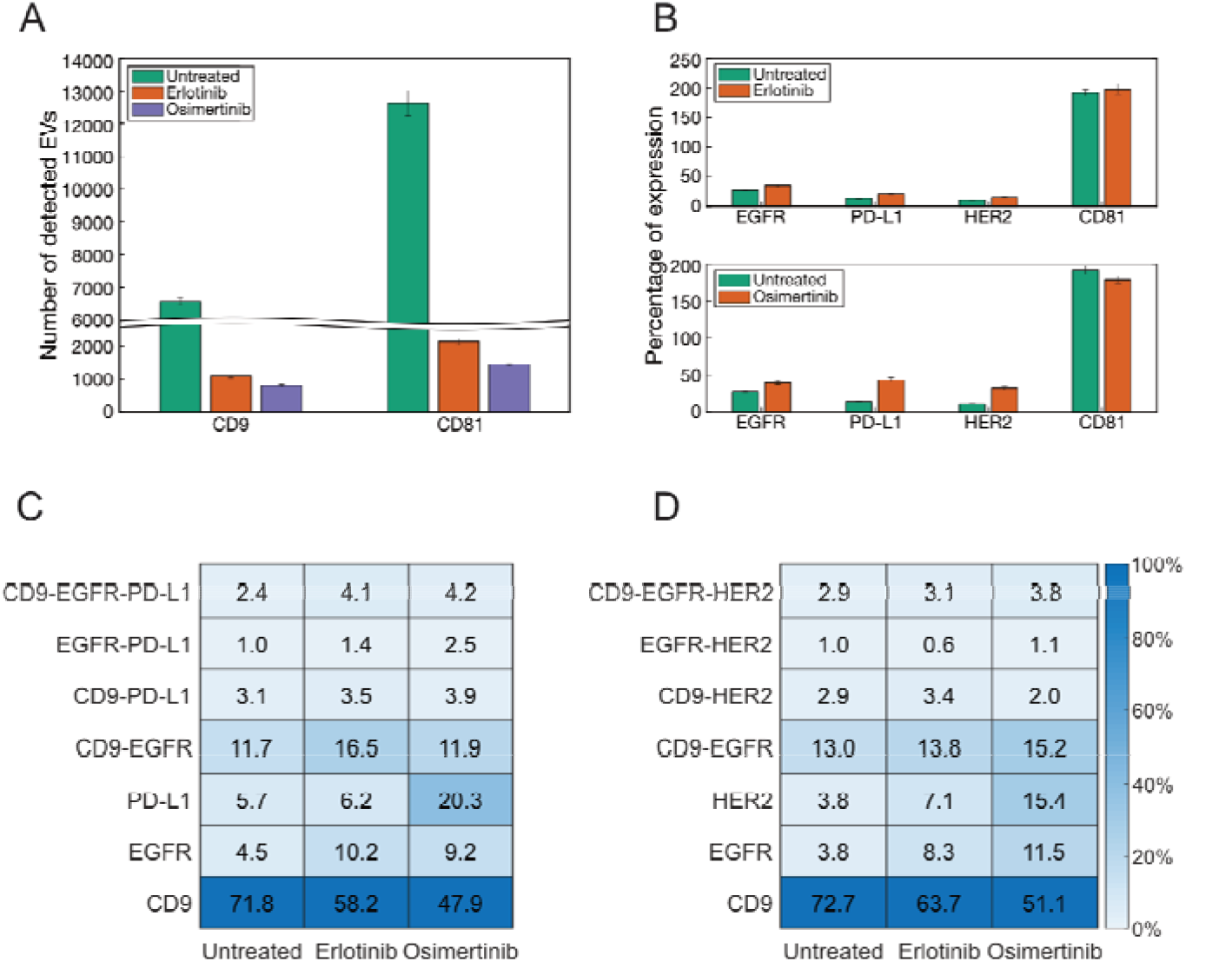
Effect of EGFR-TKI treatment on EVs membrane proteins from EGFR-mutant NSCLC cells Figure 3 Results from the fluorescent measurements on the EV samples. Number of identified CD9 and CD81 particles in the untreated, erlotinib and osimertinib treated samples (A). Number of identified particles normalized with respect to CD9 (B). Vertical bars represent standard deviations. Heatmap of colocalization between CD9, EGFR, and PD-L1 (C). Heatmap of colocalization between CD9, EGFR, and HER2 (D). Here, each row of the heatmaps corresponds to EVs positive ONLY to the mentioned proteins. CD9 and CD9-HER2 are therefore two disjoint sets. Percentage is in relation to total distinct identified EVs.

For CD9 and EGFR, the two proteins analyzed multiple times in the experiment, the average count was used. We found that all the proteins were expressed in all samples, albeit with different abundances. PD-L1 positive EVs is roughly three times as abundant in osimertinib compared to in the untreated sample; 44% ± 1.4% vs 14% ± 0.5%. For erlotinib the percentage of EVs positive for PD-L1 is closer to the untreated (20% ± 1.4%). HER2-positive vesicles follow a similar trend with the protein being slightly less than three times as common in the osimertinib sample (32% ± 2.3%) compared to untreated one (11% ± 0.3%). Once again, the percentage of HER2 positive EVs in the erlotinib sample (15.3% ± 1.4%) is closer to the untreated sample. EGFR is increased in both treated samples compared to the untreated. For osimertinib, this increase is almost 1.5-fold (comparing osimertinib expression of 40% ± 2.9% with the untreated level of 28% ± 1.1%). Erlotinib expresses a lower percentage of EGFR than osimertinib (35% ± 2.1%), but still higher than untreated.

Next, we analyzed the colocalization of proteins in the different samples. The results are presented in Figure 3C-D. In the experiment colocalization CD9, EGFR, and PD-L1, it is clear that CD9 favors colocalization with EGFR compared to PD-L1 for all samples (Figure 3C). We did not observe any major difference in the co-localization of EGFR with CD9 after EGFR-TKI treatment although the proportion of EVs only expressing EGFR was increased after either of the EGFR-TKIs. There was a clear difference in PD-L1 positivity after osimertinib treatment relative to the either untreated or erlotinib treatment however, PD-L1 was not evidently co-localized with CD9 or EGFR. HER2 positive EVs also indicated a similar trend (Figure 3D). Despite the high count of CD81 identified in the experiments, we could not establish anything noteworthy from colocalization of CD9, EGFR, and CD81.

Because of the overexpression compared to the other markers, the largest subpopulation was EVs only positive for CD81. The heatmap is shown in Figure S5. It should, however, be noted that the five proteins we studied here is only a small fraction of the hundreds of surface proteins identified in EVs derived from the H1975 cell line [28].

The fluorescence intensity distributions detected from individual EVs are presented in Figure 4. The intensity in this case represents a semi-quantitative measure of the copy number of membrane proteins, i.e., the expression level per EV. The distribution of expression levels on individual EVs for all the markers except CD9 is presented in the figure. This is because the EGFR-TKI treatment did not influence CD9 expression and the profiles for the integrated intensity of the CD9 particles remains similar prior and post EGFR-TKI treatment (p > .05), as shown in Figure S6. Although the intensity profile for CD81 drops in median value when comparing the untreated median (56503 i.u.) with the treated samples (erlotinib: 47346 i.u.; osimertinib: 38691 i.u.) (p < .001), the shape remains similar across the samples with a high population density in the lower intensity region. Such a single mode distribution of tetraspanins have been shown in other cell lines [26]. One could envision that EGFR-TKI treatment would influence EGFR expression on the cells in either direction for erlotinib or osimertinib and as EVs reflects their parental cell of origin that should also be reflected in EVs. Indeed, we found that EGFR undergoes a change in the distribution between the EVs isolated from untreated and EGFR-TKI treated cells. In EVs from the untreated sample (median: 27105 i.u.), two intensity peaks can be seen: one around 10^4^ i.u. and the other around 3 × 10^4^ i.u. For the EVs isolated from EGFR-TKI treated cells, the higher peak in the EGFR profile has flattened out and the lower intensity peak has increased, see Figure S7. Their medians are 20478 i.u. for the erlotinib-treated sample and 15062 i.u. for osimertinib-treated sample. This change is most distinct in the EVs obtained after osimertinib treatment, but the pattern is also visible in the erlotinib-treated sample.

**Figure 4.**
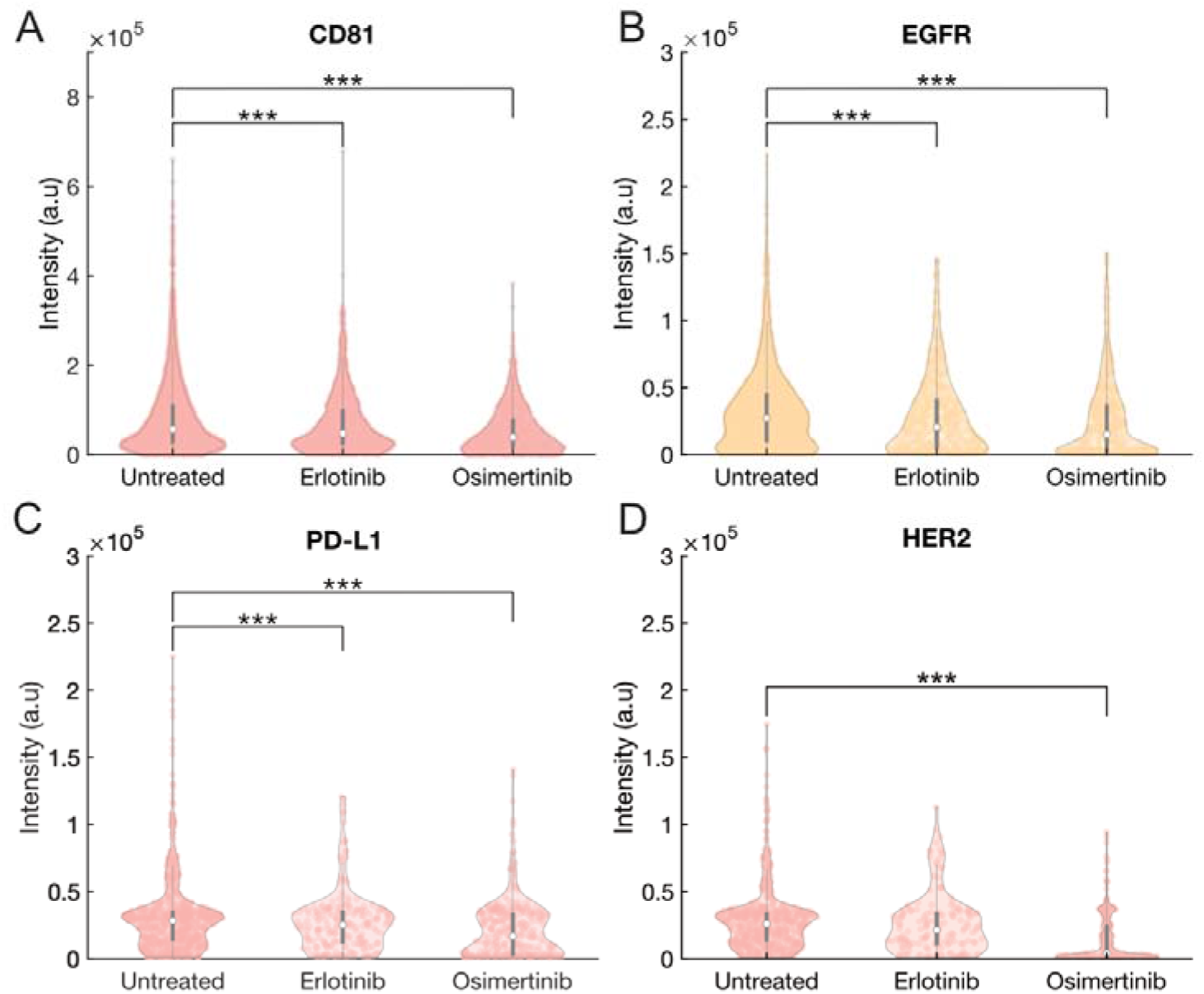
Violin plots of the fluorescence intensity distributions of the proteins. Violin plots of the fluorescence intensity distribution of the CD81 (A), EGFR (B), PD-L1 (C), and HER2 (D) particles. White circle indicates median, width indicates frequency, and grey line indicates first-third quantile. Decrease in intensity in the treated samples compared to the untreated sample with *** p <.001 by Student’s t-test.

*EGFR*-mutant NSCLC cases are reported to “immune cold” relative to other cases because of multiple alterations in the tumor cells per se e.g., increased PD-L1 expression but also because of changes in the tumor microenvironment [19]. PD-L1 is also reported in EVs from NSCLC cells including those with EGFR-mutations and as a next step in our analyses we therefore studied heterogeneity in PD-L1 expression in EVs isolated from cell culture media of H1975 cells prior and post EGFR-TKI treatment [18]. We observed two peaks in PD-L1, one of proteins with intensity between 2 × 10^4^ i.u. and 3 × 10^4^ i.u., and a smaller one around 1 × 10^4^ i.u. In context of alteration after EGFR-TKI treatment, the intensity distribution is spread out for EVs isolated after erlotinib treatment while the density of the two peaks are swapped for EVs obtained post osimertinib treatment. The decrease in the median intensity from untreated (28310 i.u.) to the treated samples (erlotinib: 25154 i.u.; osimertinib: 16770 i.u.) is statistically verified (p < .001). Thus, our results indicate that EVs from *EGFR*-mutant NSCLC cells are heterogenous in their PD-L1 expression level and that EGFR-TKI treatment impact PD-L1 expression level in EVs. Given that PD-L1 in EVs are reported to influence the cytotoxic effect of immune cells vs tumors in particular cytotoxic T-cells in other tumor types e.g., malignant melanoma, our results may have implications also for the response to immune checkpoint inhibitors in NSCLC [29].

There is evidence that HER2 is a bypass driver in 1^st^ generation EGFR-TKI as ablation of HER2 function is reported to influence the response to 1^st^ generation EGFR-TKI in *EGFR*-mutant NSCLC [30], [31]. In contrast, the 3^rd^ generation osimertinib is reported to be effective also in NSCLC cells/tumors that have increased HER2 expression [32]. We therefore analyzed the heterogeneity in HER2 expression in EVs prior and post EGFR-TKI treatments. Indeed, our study reveals an interesting change in profile for HER2 in single EVs prior and post treatment. In EVs from untreated or erlotinib treated H1975 cells, HER2 had a one more prominent peak at higher intensity and a smaller peak at lower intensity for the untreated sample which is being spread out in EVs obtained after erlotinib treatment. However, the HER2 profile for EVs obtained after osimertinib-treatment is concentrated around lower intensity levels, while a small peak about 4 × 10^4^ i.u. also exists. The decrease in median intensity from untreated (25944 i.u.) to osimertinib (4080 i.u.) treated sample was statistically significant (p < .001). This is a larger decrease compared to the median of the erlotinib sample (21358 i.u.), which could not be statistically verified (p > .05).

### Cisplatin does not change level of expression of EGFR and PD-L1 on H1975 EVs

In order to further investigate if the observed differences on the EV membrane proteins were specific to EGFR-TKIs, a separate set of experiments were done with the same cell line but treated with cisplatin which is a non-EGFR-TKI chemotherapeutic drug. Two doses of cisplatin (12.5 μM, 25 μM) was compared with untreated H1975 vesicles. The result is presented in Figure 5. The relative number of vesicles for EGFR and PD-L1, Figure 5A, after being normalized to CD9 shows no dramatic difference between treated/untreated samples.

**Figure 5.**
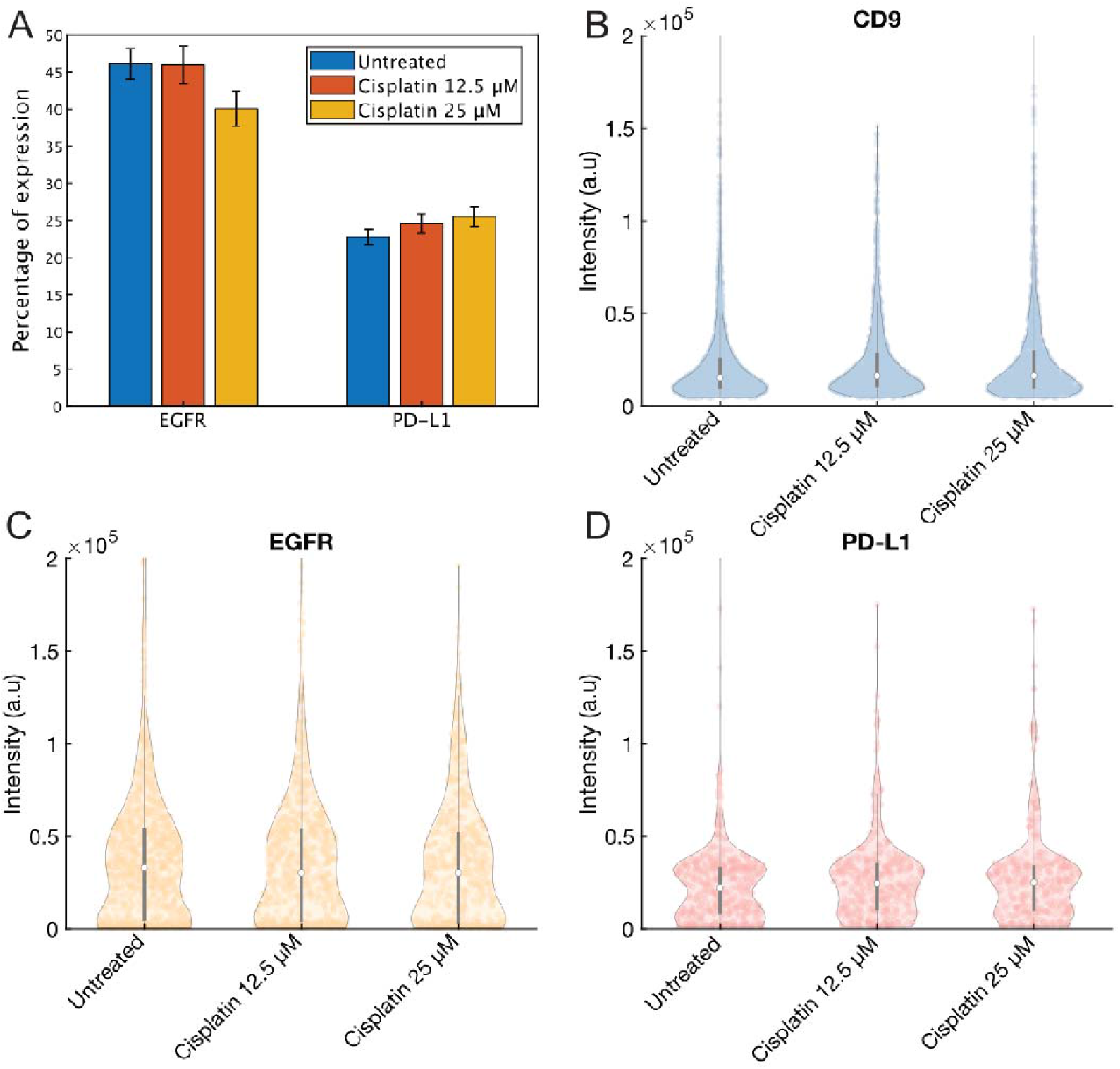
Results from the cisplatin investigation. Number of identified particles normalized with respect to CD9 (A). Violin plots of the intensity distributions of the CD9 (B), EGFR (C), PD-L1 (D) particles. White circle indicates median, width indicates frequency, and grey line indicates first-third quantile. No difference in intensity in the untreated compared to the treated samples on p¤>.05 by Student’s t-test.

The levels are roughly similar before and after treatment. The intensity distributions for CD9, Figure 5B, is also similar across the three samples. Similarly, the intensity of EGFR (Figure 5C) also did not changed after treatment. Finally, the intensity distributions of PD-L1 are shown in Figure 5D. Once again, the two intensity peaks are abundant in all three samples. No change is seen comparing untreated and treated samples. None of the three proteins showed a decrease of intensity on a statistical level (p >.05).

## Discussion

Precision cancer medicine for metastatic NSCLC have changed the treatment regimen and accordingly their outcome. For *EGFR*-mutant cases, the introduction of EGFR-TKIs have improved the survival rate [33]. Yet responses are often heterogenous and development of resistance is common even with 3^rd^ generation osimertinib, which are currently being explored as adjuvant treatment for early NSCLC cases [9], [34]. As a result, there is a need for non-invasive diagnostic and treatment monitoring approaches in order to detect the indication of such resistances at their early manifestation. This will allow the clinician to select a more appropriate treatment course. In this context, circulating EVs offer unique advantages given their high abundance in plasma/serum as well as their rich molecular content. Unlike tissue biopsies, EV-based liquid biopsies can be frequently collected and examined, allowing one to monitor the temporal evaluation and thereby offering a more accurate prognosis. Secondly, EVs containing both proteins and nucleic acids from their parental cells, allows one to follow signaling cascades both at protein and RNA level, and hence increases the diagnostic accuracy. Finally, the content of EVs is better suited to capture the tumor heterogeneity. However, despite of these benefits, using EVs as a source of tumor biomarker has proven to be extremely challenging due to the high heterogeneity in their phenotypes even when they are collected from a single cell type. The recent development of single EV technologies has, therefore, brought in a renewed sense of optimism. The focus of the present work is to explore the prospect of using EVs as source of biomarker for treatment follow up in the case of EGFR-TKI treatment.

We have earlier reported on the cell viability among the two TKIs 1^st^ generation erlotinib and the 3^rd^ generation osimertinib using the H1975 cell line [24]. Out of the two treatments, the latter one induces the strongest effect on the cell viability. The cell counts reduce to 50% after 72 hours of treatment and with a small dosage (0.1 μM). In comparison, the treatment with erlotinib induces a negligible effect under the identical treatment condition. The key question that we are trying to address here is if there is any signature on the membrane proteins of the EVs, released by these cells under different treatment, that correlates with the observation made on the cell viability. In other words, when a tumor is responsive or refractory under such PCM treatments, how do the tumor cells respond in terms of the membrane protein composition of EVs released by them. Although, EVs are known to carry a large number of different membrane protein, we selected five membrane proteins in this study. This includes the tetraspanin CD9 and CD81 which are common EV markers alongside EGFR; the bypass resistance driver HER2 and PD-L1, which is an important target for immune check point inhibitor treatments. The selection is also motivated by their important roles in the context of EGFR-TKIs as explained earlier. Our focus in the present study was to investigate, i) how the population of EVs expressing these membrane proteins and their combinations changes upon various treatments? and ii) how do the expression levels of these protein on single EVs and their distribution in the sample change? To make a further distinction between EGFR-TKIs and chemotherapy medication used to kill proliferating cells, the present study also includes EVs derived from cells those are treated with cisplatin with different dosage.

### i. Changes in EV population bearing the selected membrane proteins

As presented in Figure 3B, the population of EGFR, PD-L1 and HER2 positive EVs clearly increases upon osimertinib treatment. Among the selected markers, PD-L1 positive EV population shows the largest increase (about 3-fold). Interestingly, cisplatin treatment which induces even stronger cell death does not apparently have similar effect on the EGFR or PD-L1 positive EV population (Figure 5A). The co-expression analysis presented in Figure 3C and D shows some additional features. It appears from the heatmaps that the population of EVs expressing more than one of the selected proteins per EV, remains quite the same irrespective of the treatment types. It is only those EV population expressing either one of the three proteins (PD-L1/EGFR/HER2) that exhibits distinct increase in the population upon the treatment. In summary, the above analyses indicate that only a subset of PD-L1/EGFR/HER2-positive EV population is increased because of the TKI treatments. Within the same subsets, osimertinib treatment significantly increases the PD-L1/HER2 positive EV populations as compared to the erlotinib treatment.

### ii. Changes in the expression levels on single EVs

In addition to their population, the selected markers also show interesting differences in their expression levels on individual EVs with respect to the treatments. As presented in Figure 4A-D and 5B-D, the tetraspanins expression show a unimodal distribution. However, the EGFR/PD-L1/HER2 positive EV population show a bimodal distribution indicating the presence of two distinct subsets differing in their level of expression. While the distribution of the tetraspanins positive EVs remains similar upon the treatments, significant alterations was observed in the case of PD-L1/EGFR and HER2 positive EVs when treated with TKIs. In the case of osimertinib treatment, the mean, and the median level of expression of these markers dropped drastically with the strongest impact being on the subset having higher intensity (Figure 4B-D). In comparison, erlotinib treatment retains similar modal distribution of the untreated sample, although shows a slight decrease in the overall intensity level. Cisplatin treatment, on the other hand, shows no alteration to the intensity distribution of the analyzed markers.

Clearly, the single-EV analysis of the TKI-treated H1975 cells reveals many interesting features which have not been captured earlier, although some indications of the observed differences have been previously reported in primary cells and in bulk EV analysis. This includes indications of a general decrease of CD81, EGFR, and PD-L1 in EVs post treatment [35]. However, the analysis is based on the total protein analysis and not calculated per vesicle, as is shown in this study. Wu et al published a study in 2021 proving a connection between wild-type EGFR in exosomes and osimertinib resistance in H1975 cells [36]. In the study, it is concluded that osimertinib triggered exosome releasing in cells which EGFR-mutated cells pick up causing osimertinib resistance. This could explain the increase of EGFR we see in our osimertinib sample compared to the untreated sample. HER2 is reported to be a bypass-driver for 1^st^ generation EGFR-TKI while for osimertinib, HER2 overexpression doesn’t block response at least not in NSCLC cultured *in vitro* [30], [31], [32]. This study is novel in discovering how the HER2 expression changes post EGFR-TKI treatment on vesicular level. Upon literature survey it is clear that the EV studies on EGFR-TKI treatments are scarce, and on single EV-level even more.

Increased PD-L1 expression in osimertinib or erlotinib-resistant NSCLC cells has previously been documented [18] [37]. PD-L1 expression have earlier been reported in EVs from EGFR-mutant cells however to our knowledge, the alteration after EGFR-TKI treatment has not been reported. We also observed interesting alterations on PD-L1 intensity in EVs prior and post EGFR-TKI treatment. While EVs from the untreated sample has two intensity peaks and favors the higher, the EVs obtained after osimertinib treatment favors the low-intensity peak. Our interpretation is that these two peaks could correspond to subpopulations of EVs with lower and higher abundance of the marker. By combining the results from the numerical data with the intensities it seems like the EVs isolated post EGFR-TKI treatment are more positive to PD-L1, but with reduced number of proteins per vesicle. As it has been demonstrated in other tumor types that PD-L1 expressing EVs may block effect of ICI treatment it is interesting to note that PD-L1 alterations occur on EVs after EGFR-TKI which may indeed contribute to the observed immune cold phenotype of EGFR-mutant NSCLC cases [29], [19], [38], [39], [40].

As for the intensity profiles of EVs for the proteins, we see a decrease in the intensity of the EGFR expressing EVs for the samples coming from EGFR-TKI treated NSCLC cells vs. untreated ones. From a treatment perspective this could be explained by the fact that NSCLC cells with high EGFR level are most vulnerable to EGFR-TKI treatment and are killed off early on and hence those with lower EGFR level will be the main source of EVs. However, we didn’t see any clear difference between erlotinib and osimertinib in this context which is puzzling as these TKIs have different EGFR-TKI sensitivity in the studied NSCLC cell line [21], [20], [24]. However, the experiments on the two cisplatin samples showed no decrease of either EGFR positive EVs or mean intensity. This is expected, as cisplatin is not an EGFR specific treatment. Nevertheless, results show that our analytical method for EGFR can reveal alterations after EGFR-TKI treatment and a next step would be to apply this method on EVs from plasma of EGFR-TKI treated NSCLC patients with or without response to given EGFR-TKI.

From a biological and clinical perspective our study show feasibility to address the heterogeneity of EV surface proteins taking both number of EVs which express certain marker as well as their diverse intensity. We demonstrate this for EGFR, HER2 and PD-L1 which all are relevant in context of EGFR-TKI treatment of NSCLC. The next step would be to further apply the method in EVs isolated from plasma/serum of EGFR-TKI treated NSCLC where bulk analyses have revealed candidate markers linked to response. We also foresee that our method should be combined with analyses of exoRNA mutations and/or cf-DNA to cover different aspects of EGFR-TKI resistance.

## Conclusions

In summary, we demonstrated here that a single-EV analysis of membrane proteins can reveal the response of a tumor which is under a treatment such as EGFR-TKIs, providing clinician an early indication of e.g., a possible drug resistance. The study was performed with an EGFR-mutant NSCLC cell line and three different treatment types; 1^st^ generation EGFR-TKI erlotinib, 3^rd^ generation EGFR-TKI osimertinib and a chemotherapeutic drug, cisplatin. The chosen cell-line was resistant to erlotinib and responsive to osimertinib and cisplatin, as also verified by a cell viability study. Thus, the model system allowed us to compare the differences in EV signatures in the different response types. In contrast to the erlotinib treatment, the osimertinib-treated samples showed a significant increase in the population of PD-L1, EGFR, and HER2. The signature was dominant for the case of PD-L1 expressing EVs. Interestingly, the effect was mostly influencing a subset of EVs expressing these proteins, e.g., the EV-subset that were only positive to HER2 or PD-L1, and not coexpressing the other three proteins studied here. Furthermore, the two different EGFR-TKI treatments also had a distinctive alteration on the distribution of expression levels on single EVs for the marker PD-L1, EGFR and HER2. None of the observed differences induced by the EGFR-TKIs could be reproduced with the chemotherapeutic drug. Thus, the work presents a clear evidence that single-EV profile carry distinct signatures which are indicative of tumor response to EGFR-TKIs. If validated on a clinical cohort, such a method can help to derive valuable diagnostic and prognostic signatures from liquid biopsy analysis of cancer patients.

## ACKNOWLEDGEMENTS

This project was supported by a generous grant from the Erling Persson Family Foundation. A.D., F.S., and S.S.S. acknowledge also the grant funded by the Swedish Research Council (Contract No. 2016–05051). The biological part of the study was granted to R.L, K.V and P.H from the Swedish Cancer Society (contract no. CAN 2018/597 and CAN2021/1469 Pj01 H), the Stockholm Cancer Society (contract no. 201202 and 191293) and Stockholm County Council (909121). The project was also supported by a cooperation grant from Stockholm County Council to the entire group contract. no. FoUI-966345. The Karolinska team would also like to thank MSc Vasiliki Arapi and Dr Vitaliy Kaminskyy for helping with some EVs isolations.

SELA is supported by the Swedish Research Council and SSF-IRC (FormulaEx). Figure 1 was realized partly using BioRender.com.

## DECLARATION OF INTEREST STATEMENT

No competing interests need to be declared by the authors.

## Notes

### Competing Interest Statement

The authors have declared no competing interest.

