## Supplemental information for "Analyses of single extracellular vesicles from non-small lung cancer cells to reveal effects by Epidermal growth factor inhibitor treatments"

Figure S1 Cell morphology before harvesting EVs for the different samples. The cells were treated 48h prior imaging. Number of counted cells: Untreated: 6083333, osimertinib (0.1 µM) 3600000, cisplatin (12.5 µM) 2666667, and cisplatin (25 µM): 1466667.

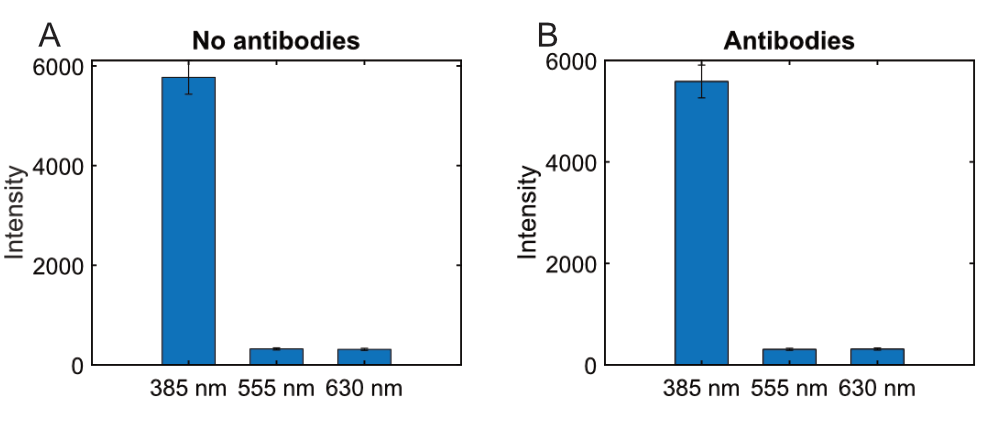

Figure S2 Backgrounds intensities of images with PLL as capture agent without (A) and with (B) antibodies. Bars represent standard deviation.

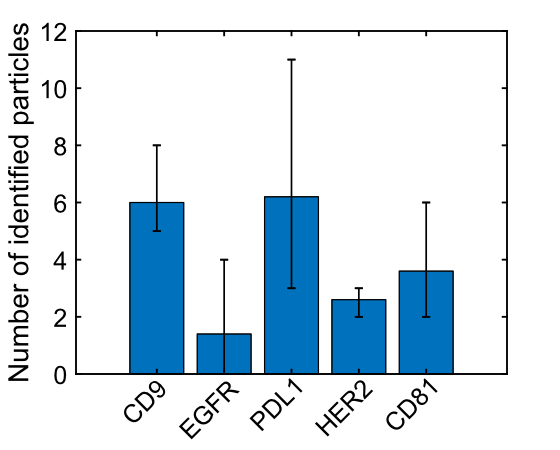

Figure S3 Number of identified particles on a functionalized without any EVs but with antibodies. The reported number is the average of 5 images. Bars represent lowest/highest reported number of particles in an analyzed image.

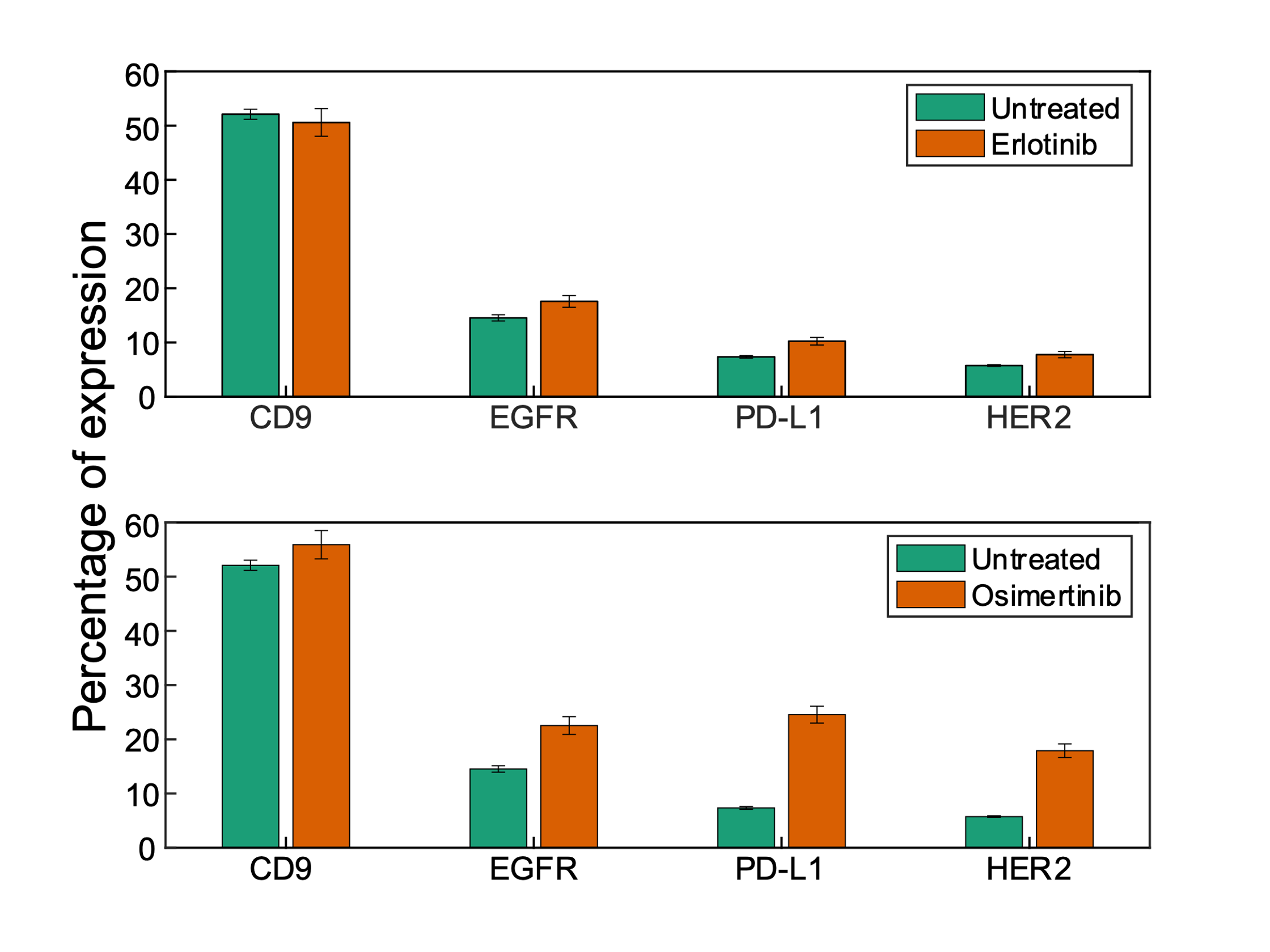

Figure S4 Percentage of identified particles. Each group corresponds to particles positive for a specific antibody. The colors represent untreated, erlotinib-, and osimertinib-treated samples respectively. All samples have been normalized with respect to CD81. Vertical bars represent standard deviation.

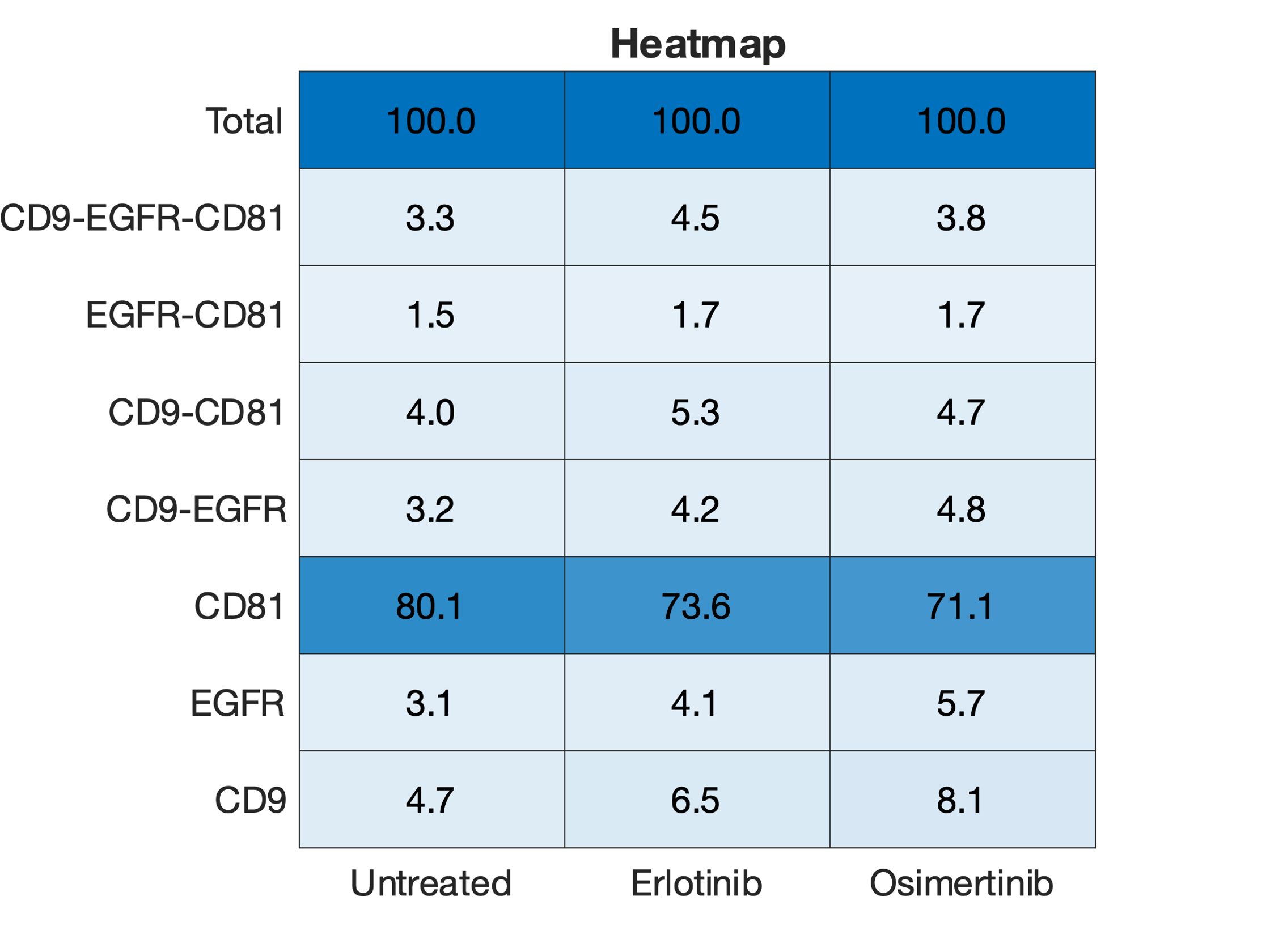

Figure S5 Heatmap of colocalization between CD9, EGFR, and CD81.

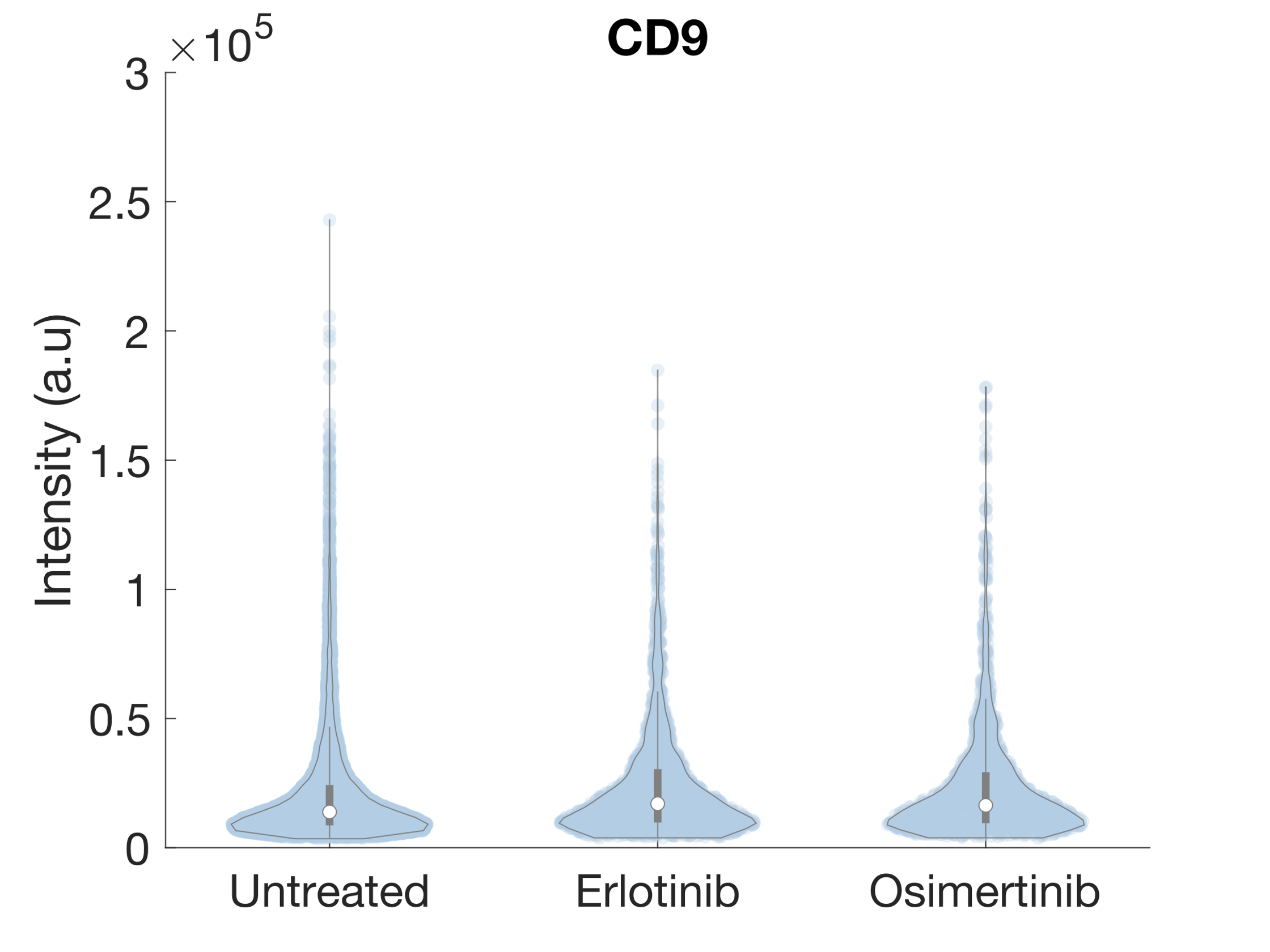

Figure S6 Violin plot of the fluorescence intensity distributions of the CD9 particles.

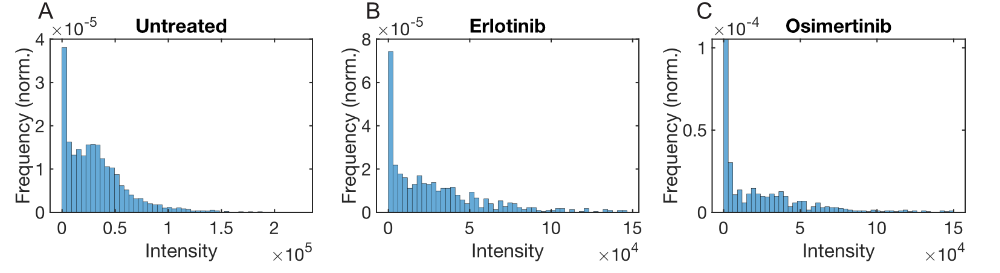

Figure S7 normalized intensity frequency of the identified EGFR particles for the untreated (A), erlotinib-treated (B) and osimertinib-treated (C) sample.
